# Efficient neural codes naturally emerge through gradient descent learning

**DOI:** 10.1101/2022.05.11.491548

**Authors:** Ari S. Benjamin, Ling-Qi Zhang, Cheng Qiu, Alan Stocker, Konrad P. Kording

**Affiliations:** Department of Bioengineering, University of Pennsylvania, Philadelphia, Pennsylvania, USA; Department of Psychology, University of Pennsylvania, Philadelphia, Pennsylvania, USA; Department of Neuroscience, University of Pennsylvania, Philadelphia, Pennsylvania, USA

## Abstract

Animal sensory systems are more sensitive to common features in the environment than uncommon features. For example, small deviations from the more frequently encountered horizontal orientations can be more easily detected than small deviations from the less frequent diagonal ones. Here we find that artificial neural networks trained to recognize objects also have patterns of sensitivity that match the statistics of features in images. To interpret these findings, we show mathematically that learning with gradient descent in deep neural networks preferentially creates representations that are more sensitive to common features, a hallmark of efficient coding. This result suggests that efficient coding naturally emerges from gradient-like learning on natural stimuli.

## Introduction

Careful psychophysical studies of perception has revealed that neural representations do not encode all aspects of stimuli with equal sensitivity (Fechner, 1948). The ability to detect a small change in a stimulus, for instance, depends systematically on stimulus value. A classic example of this is the so-called ‘oblique effect’ in which changes in visual orientation are easier to detect near vertical or horizontal than oblique orientations (Appelle, 1972). The fact that these sensitivity patterns are ubiquitous and widely shared between animals motivates us to study the potential underlying reasons why they exist.

The efficient coding hypothesis has become a standard explanation for the emergence of these non-homogeneous sensitivity patterns (Barlow, 1961). It predicts that sensory systems should preferentially encode more common aspects of the world at the expense of less common aspects, as this is the most efficient way (in the information-theoretical sense) to make use of limited coding resources. Indeed, perceptual sensitivity typically reflects the statistics of the visual environment (Coppola et al, 1998; Ganguli and Simoncelli, 2010; Girshick et al, 2011; Wei and Stocker, 2015, 2017). While much is known about efficient neural codes and their link to the stimulus statistics and perceptual behavior, the mechanisms that give rise to such codes remain unknown.

Brains are not born with fully developed sensory representations. Many developmental studies of perception in infants and young children have shown that visual sensitivities improve with age and visual experience even until adolescence (Armstrong et al, 2009; Braddick and Atkinson, 2011; Mayer and Dobson, 1982; Teller and Movshon, 1986). The maturing optics of the developing eye and retina explain some of this improvement, but much of the improvement depends on visual experience and is due to neural changes downstream (Banks and Crowell, 1993; Maurer et al, 1999; Movshon and Kiorpes, 1993; Gold et al, 1999; Schoups et al, 2001). This suggests that perceptual sensitivity depends on the neural representation of sensory information and how these representations change with experience during development.

We hypothesize that general, task-oriented learning is the mechanism that gives rise to efficient sensory representations in the brain. Any gradual learning process can only learn so much at a time. This means that an effective learning algorithm should prioritize learning more important aspects before less important ones. Conveniently, features that are more common are also easier to learn from a limited exposure to the world. This suggests that effective learning rules for neural networks might naturally produce better representations for common features, hence providing a mechanism for the emergence of efficient codes. This phenomenon is equivalent to the notion that learning provides a second, implicit constraint on neural coding in addition to the explicit constraint imposed by the limited neural resources. Our hypothesis directly predicts that we should find forms of efficient coding not just in biological neural networks but also in other learning systems. Especially, we expect artificial neural networks trained to perform visual recognition tasks to exhibit efficient neural representations similar to those found in the visual cortex of biological brains despite their many differences in their local structural properties (e.g. noise) and connectivity.

A study of the consequences of effective yet gradual learning on sensory representations must begin from a specific learning rule. One canonical learning rule is gradient descent, which proposes that neural updates are as small as possible to elicit a given improvement in behavior. Though the brain may use more complicated learning rules, gradient descent is arguably the simplest rule for general learning and thus a baseline for theorizing about learning in the brain. If gradient descent produces efficient codes, this would provide a strong proof of principle that efficient codes can emerge from general-purpose learning algorithms.

To show that efficient coding emerges from gradient descent requires a formal understanding of how learning with gradient descent biases what is represented about the stimulus. This parallels an active effort in the study of deep learning. It is now recognized that what neural network learn about their inputs is constrained not just by their connectivity but also implicitly by their learning algorithm (Zhang et al, 2021). Many potential implicit constraints have been propose due to their importance in explaining why large neural networks work well on unseen data (i.e. generalize) (Jacot et al, 2018; Neyshabur et al, 2017; Smith and Le, 2017; Tishby and Zaslavsky, 2015). One prominent theory is gradient descent in a multilayer network produces nontrivial learning dynamics that supplies key biases on what is learned first (Arora et al, 2019; Gidel et al, 2019; Gunasekar et al, 2018; Razin and Cohen, 2020; Saxe et al, 2013). This raises the possibility that such ideas could also demonstrate whether gradient descent learning is biased towards efficient codes.

In this paper, we demonstrate that learning with gradient descent biases feature learning towards common input features, thus reproducing the relationship between stimulus statistics and perceptual sensitivity (Fig. 1). This effect occurs in otherwise unconstrained and noiseless networks as well as for multiple learning objectives (i.e. not limited to information maximization). We examine two model systems. First, we show that deep artificial networks trained on natural image classification show similar patterns of sensitivity as humans, and that this is a partly a consequence of image statistics (also see Benjamin et al (2019); Henderson and Serences (2021)) but is also partially due to factors inherent in network architecture. We then leverage results from the study of linear networks to mathematically describe how gradient descent naturally causes learned representations to reflect the input statistics. To demonstrate that this framework can be applied to explain development, we also show that changes in sensitivity resembling changes in visual acuity in human children can be reproduced in a simple model trained with gradient descent on natural images. Our results show how learning provides a natural mechanism for the emergence of a non-uniform sensory sensitivity that matches input statistics.

**Fig. 1.**
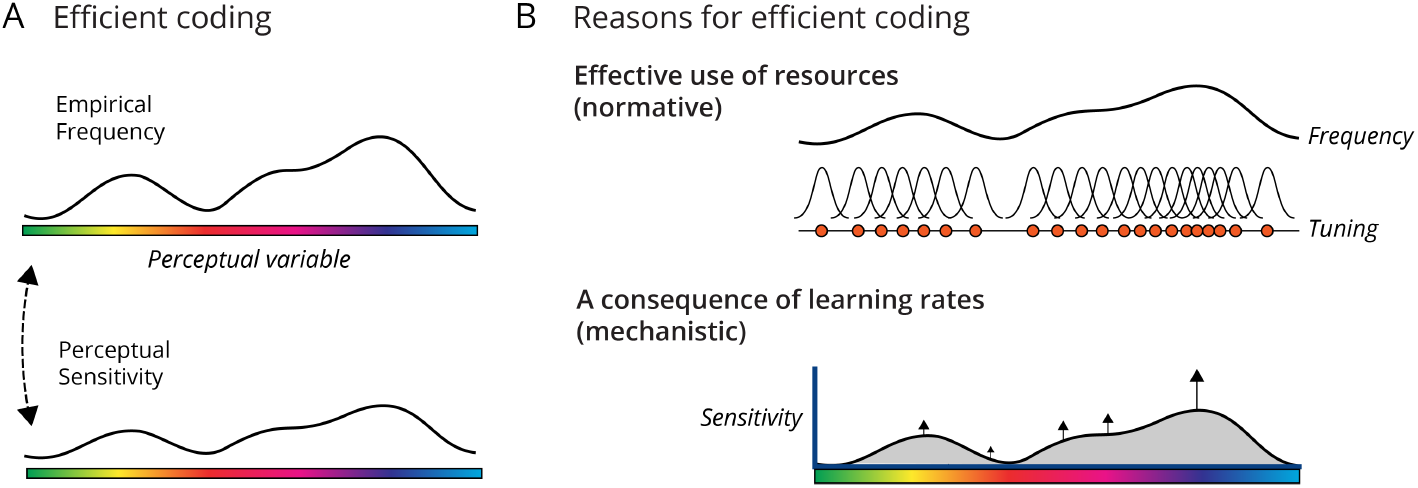
Reasons for efficient coding. A) One consequence of efficient coding is that perceptual sensitivity reflects the empirical frequency of perceptual variables. B) Efficient coding can be justified normatively as the most effective way to allocate finite neural resources to encode a stimulus ensemble. In this work we describe a mechanism for efficient coding due to learning components of the inputs at different rates dependent on their frequency.

## Results

Humans and animals show sensitivity that depends on the orientation of stimuli. In humans, the sensitivity of internal representations can be inferred from psychophysical data on discrimination thresholds (Ganguli and Simoncelli, 2010) or the empirical distribution of tuning curves in V1 (Schoups et al, 2001; Stringer et al, 2021) (Fig. 2a). In many animals, internal representations are most sensitive at near vertical and horizontal orientations (Appelle, 1972). This raises the question if neural networks trained on natural stimuli are similarly sensitive in a non-uniform way.

**Fig. 2.**
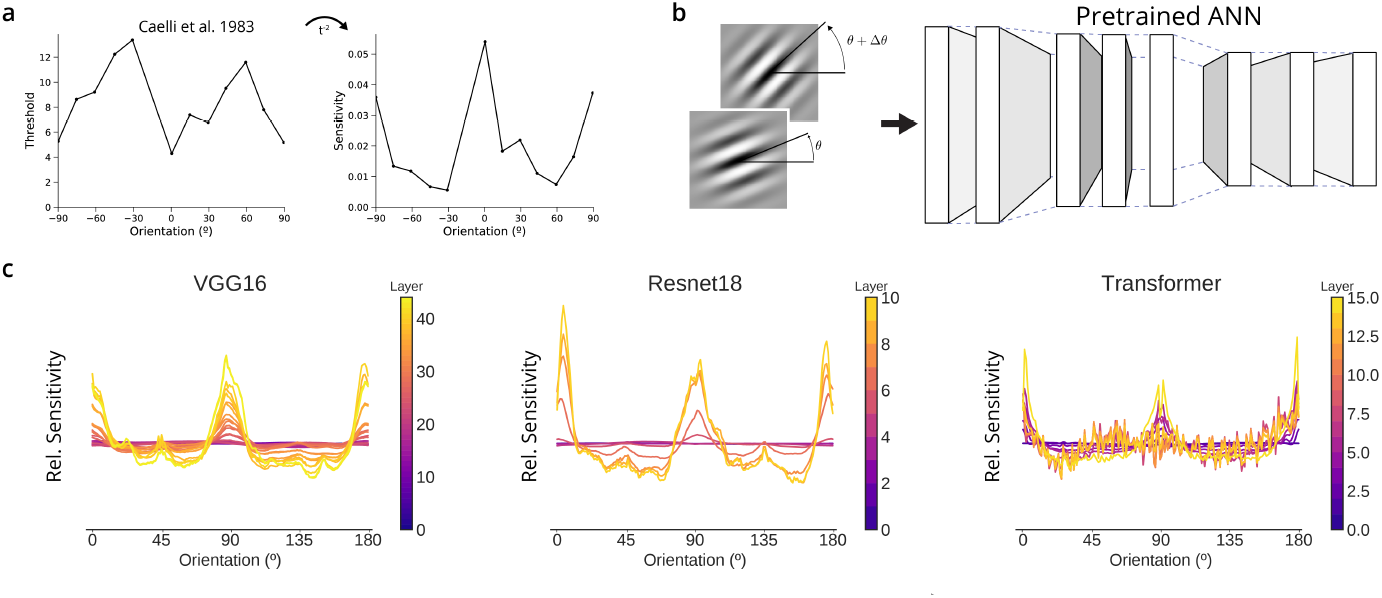
Artificial neural networks trained to classify naturalistic images show similar patterns of sensitivity as humans. a) Discrimination thresholds for orientation vary systematically in humans. The sensitivity of the underlying internal representations, as the Fisher Information, can be inferred as the inverse square of the threshold (Ganguli and Simoncelli, 2010; Wei and Stocker, 2017). b) We measured the sensitivity of each layer in an artificial network as the change in layer’s response due to a given change in orientation, i.e. the squared norm of the gradient with respect to orientation. c) Relative (normalized) sensitivity to orientation for three networks trained on ImageNet, plotted for various layers in each network.

To ask if neural networks would show a similar phenomenon, we first obtained a set of relevant networks and measured their response to artificial stimuli. We specifically investigated deep neural networks trained on the ImageNet task (Deng et al, 2009) as such networks show a number of other similarities to human ventral stream visual processing (Güçlü and van Gerven, 2015; Khaligh-Razavi and Kriegeskorte, 2014; Yamins et al, 2014). We analyzed a range of architectures, including two large convolutional neural networks (CNNs), VGG16 and Resnet18, and Vision Transformers which operate largely without convolution (Dosovitskiy et al, 2020; He et al, 2016; Simonyan and Zisserman, 2014). Then, to measure sensitivity, we measured the squared magnitude of the change in network activations given a change in the angle of oriented Gabor stimuli (Fig 2b; see Methods). For all three networks, we found that the internal representations were most sensitive to changes near cardinal orientations (Fig. 2c). The effect was more pronounced deeper in each network. The coarse pattern of sensitivity of ImageNet-trained deep networks to orientation is thus similar to that of animals and animals.

We next investigated whether this pattern was due to factors inherent in the network or due to the statistics of the inputs on which it was trained. Before training, randomly initialized networks largely do not show this pattern, nor do networks in which all weights after training are randomly shuffled (SI Fig. 1a). After training networks on a version of ImageNet in which all images are rotated by 45º, the networks lose sensitivity to cardinals and gain sensitivity to oblique angles (SI Fig. 1b). This finding recapitulates our preliminary findings and concurrent work of colleagues, and points to an origin in image statistics (Benjamin et al, 2019; Henderson and Serences, 2021). However, we also found that networks trained on rotated images do partially retain sensitivity to cardinal orientations; they do not simply rotate their sensitivity by 45º (SI Fig. 1b). This indicates that increased sensitivity to cardinals is partially due to factors inherent in the convolutional architecture. Indeed, we found that the use of spatial pooling with overlapping receptive fields (such as in AlexNet; Krizhevsky et al (2012)) involves oversampling a rectangular grid and that this produces a significant cardinal sensitivity (SI Fig 1c). The pattern of orientation sensitivity is thus both a product of the input statistics and inherent factors like architecture.

To separate effects related to architecture and learning, we next examined the sensitivity of artificial neural networks to changes in hue, as this is unlikely to be affected by rectangular convolutional processing. We found that hue sensitivity after training was related to the empirical frequency of hues in ImageNet (Fig. 3c) measured in HSV color space. The location of the peaks of network sensitivity roughly matched the patterns of human sensitivity to changes in hues in the HSV color space (Fig. 3b). To test if this pattern is causally related to the input statistics, we trained a Resnet18 network on a version of ImageNet in which the hue of all pixels was shifted by 90º and observed a corresponding shift in the hue sensitivity (Fig. 3d). This suggests that in general the frequency of low-level visual features determines the sensitivity of artificial neural networks trained on object classification.

**Fig. 3.**
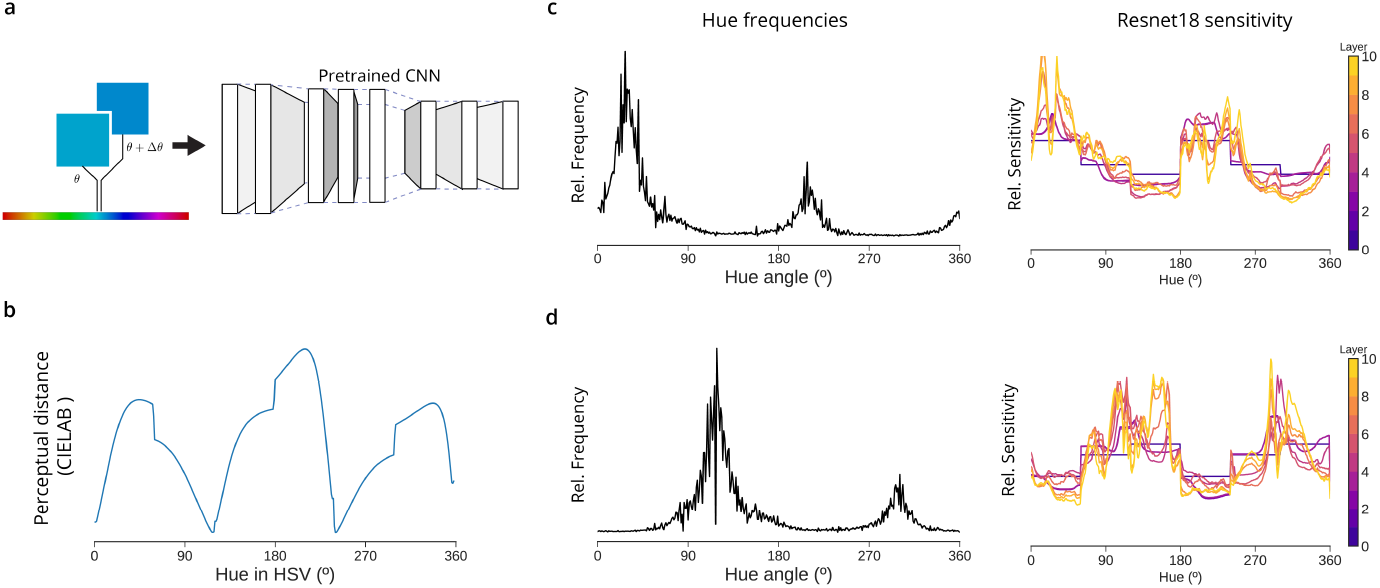
The sensitivity of ANNs to hue also matches image statistics. a) The color of a uniform image is varied; in HSV color space, saturation and value are held fixed and the hue is varied. Results are averaged over possible saturations and values. b) In humans, the sensitivity to the H axis can be inferred by the perceptual distance between uniformly spaced H values (calculated using the approximately perceptually uniform color space CIELAB) at S=V=1. c) The sensitivity to hue in each layer in a trained ResNet18 tracks the empirical frequency of hues in the ImageNet dataset. d) Training ResNet18 on a version of ImageNet in which hues are rotated results in a corresponding shift in hue sensitivity.

Sensitivity may track input statistics in artificial networks for a similar reason as in humans. In psychophysics, one leading explanation proposes that there is some constraint that limits the amount of information a neural population can contain about its inputs. Due to this constraint, an optimal code will allocate more resources (and be more sensitive) to inputs that occur frequently (Ganguli and Simoncelli, 2010; Wei and Stocker, 2016). However, the networks above are overparameterized, in the sense that internal layers contain a greater number of nodes than there are input pixels, and furthermore contain no source of information-limiting noise during evaluation. Thus, the frequency/sensitivity correspondence in artificial networks likely does not arise from an optimal encoding of the inputs despite inherent and unresolvable architectural constraints.

One alternative possibility to an explicit constraint like noise is that incremental learning via gradient descent naturally leads to a frequency/sensitivity correspondence. This hypothesis relates to the idea from connectionist models of development that the most general aspects of a problem are often learned first (Munakata and McClelland, 2003). To investigate this possibility, we analyzed a category of artificial neural networks amenable to mathematical study: deep linear networks. Deep linear networks contain no nonlinearities and are equivalent to sequential matrix multiplication. Despite their simplicity, deep linear networks show many of the same learning phenomena as nonlinear networks (Saxe et al, 2013; Arora et al, 2019) and humans (Lee et al, 2014; Saxe et al, 2019). Moreover, this simplified setting allows us to separate the effects of gradient descent from those of network nonlinearity.

How can we characterize learning in this linear setting? At a high level, the network becomes responsive to features earlier when those features are more common (Fig. 4). When learning ends due to finite training time, finite data, or saturating performance, there is a residual higher sensitivity for common features. We will expand on this phenomenon below. Overall, the link between learning rate and input frequency, combined with finite training time, is an additional inductive bias beyond what features are useful for the task and means that trained networks will tend to be more sensitive to frequent features.

**Fig. 4.**
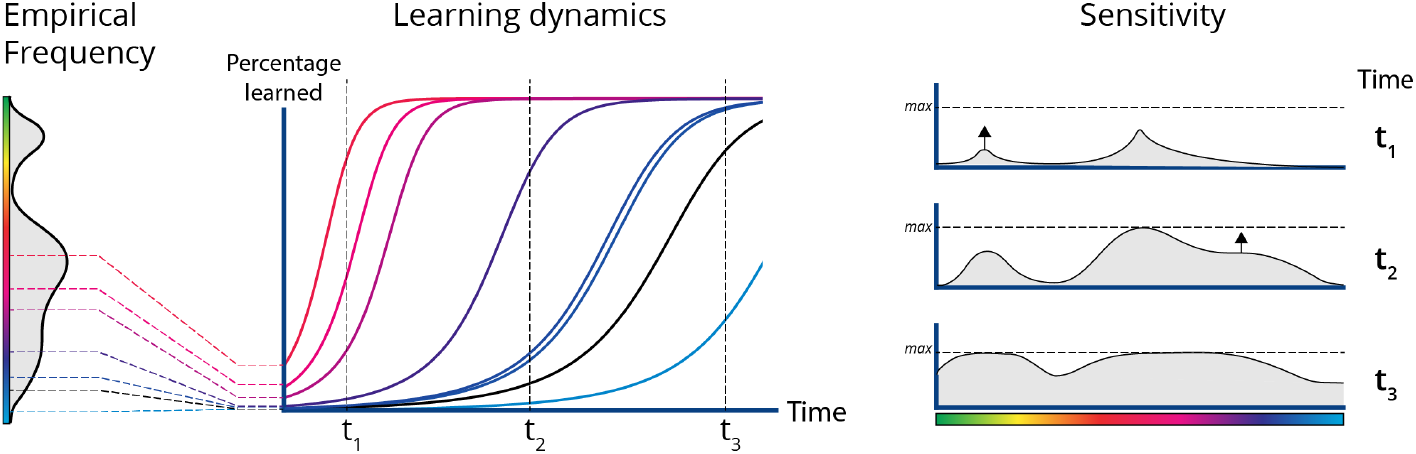
Schematic of how the learning dynamics of linear networks causes a correspondence between network sensitivity and input statistics. The learning problem is broken into components, each of which learns at a specific rate. The frequency or variance of a feature of the input data (e.g. the color red) in part determines the learning rate of the components that encode it. This means that the network becomes sensitive to frequent features first. Training may end before all features are fully encoded.

To more concretely demonstrate this phenomenon we will focus on the task of reconstructing natural images with a linear network (Fig. 5). This network can be as shallow as a single layer, in which case the reconstructed images are given by the matrix multiplication 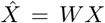. Importantly, this problem can be solved exactly with the solution that the weight matrix *W* is the identity matrix *I*. This is an unconstrained problem; if there is any non-uniformity in the sensitivity of the output 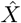 to changes in *X* it must be due to the implicit constraints posed during learning. Analyzing the output sensitivity in this simple model will help to better understand the implicit preferences of learning with gradient descent.

**Fig. 5.**
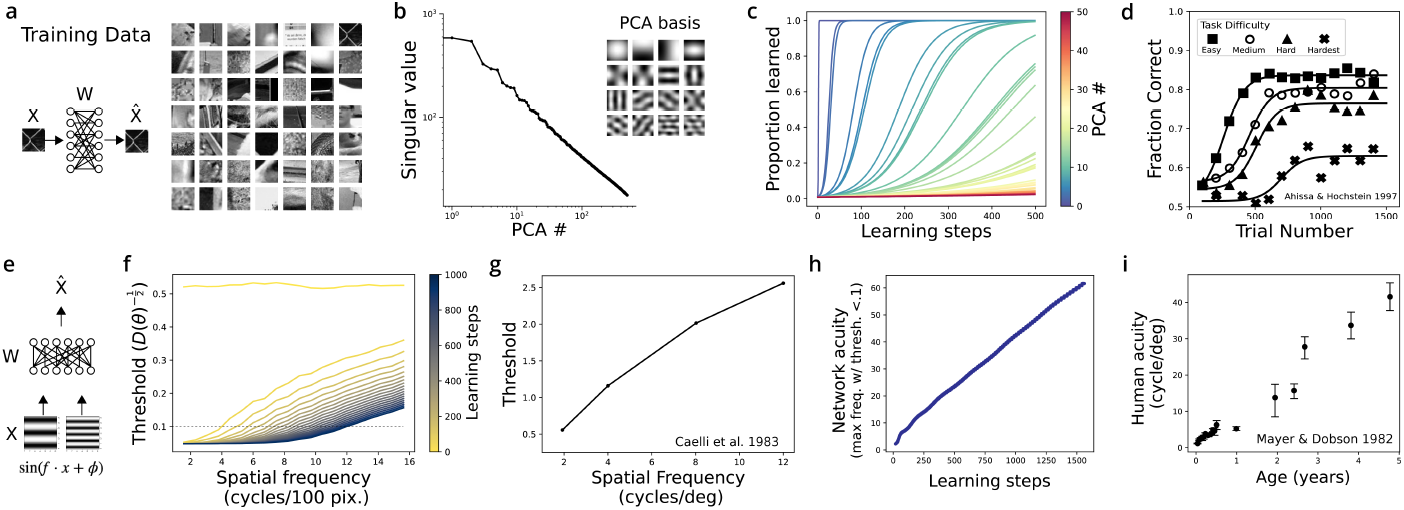
The effect of input statistics on network sensitivity can be understood with linear network models. Despite their simplicity these show human-like learning phenomena. a) We trained linear networks to reconstruct black and white patches of natural images. b) The statistics (here, variance) of each PC is given by its singular value, which for natural images shows a characteristic power law decay. c) When learning with gradient descent, the weight matrix learns each PC separately and in order of their variance. The sharpness of the sigmoidal learning curve is controlled by the network depth (SI Fig. 2) d) Human perceptual learning curves are also sigmoidal, and increasing task difficulty delays learning dynamics. Data replotted from Ahissar and Hochstein (1997); subjects trained to detect the orientation of a line, and the difficulty of the task was controlled by a masking stimulus. e-i) Sensitivity to spatial frequency. f) Every 50 learning steps we plotted the inverse square root of the sensitivity to spatial frequency, which is a proxy for detection thresholds. At each step note the linear increase above an elbow. g) Human data on spatial frequency thresholds replotted from Caelli et al (1983). h) An artificial spatial ‘acuity’ grows nearly linearly with training; ‘acuity’ is defined as the maximum spatial frequency for which the artificial threshold is below a value of 0.1. i) In infants and children, the spatial acuity – the highest spatial frequency observable for high-contrast gratings – increases linearly with age. Replotted from Mayer and Dobson (1982).

In our demonstrative task we will examine the sensitivity of the output 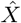 to the magnitude of each principal component that makes up an image (as provided by PCA on the inputs, Fig. 5b). The mathematical analysis for this feature is much simpler than, say, for orientation. In this case also we have some expectation as to what pattern of sensitivity the efficient coding framework predicts because the principal components (PCs) are ordered by their variance. An efficient code in the presence of independent internal noise should be more sensitive to earlier PCs. Indeed, earlier PCs are composed of lower spatial frequencies, and humans are better at detecting changes in lower spatial frequencies (Fig. 5e). If gradient descent provides a similar effect, we should find that the output 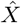 becomes sensitive to lower PCs first.

In our linear model it is possible to describe analytically how the sensitivity changes due to gradient descent. This is done by examining how the weights change. We first decompose the weights *W* via singular value decomposition (SVD), 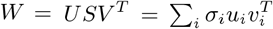, as a product of unit-length singular vectors (*u, v*) and their corresponding singular values *σ*_*i*_. The evolution of these components under gradient descent is known as long as certain basic conditions are met, such as a very small weight initialization (Arora et al, 2019; Saxe et al, 2013). One key previous finding is that the singular vectors *v* of the weight matrix rotate to align with the PCs of the input (see Theorem 2 in the Appendix) (Arora et al, 2019). Due to this alignment, the sensitivity of the output 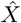 to the *i*th PC is controlled by the size of the corresponding singular value in the weights, *σ*_*i*_. This is more formally derived in Methods. If *σ*_*i*_ remains near its initialization close to zero, then the projection of data upon the *i*th PC will be filtered out and the output will not be sensitive to the corresponding PC. The sensitivity of 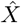 to each PC and how it changes with learning can be understood entirely by the growth of the singular values of *W*.

Thus far we have linked sensitivity to the weights, but it remains to input statistics to how the weights change. The input statistics affect the growth of the singular values of the weight matrix. Each *σ*_*i*_ grows at a different rate. For this objective of reconstruction, the growth rate of *σ*_*i*_ is proportional to the standard deviation of the corresponding *i*th PC in the data (see Methods). These decay as a power law for natural images and are shown in the spectrum in Fig. 5b. As a result, the network output will become sensitive first to the first (largest-variance) PCs and later to the later PCs. This is verified empirically in Fig. 5c. Only at infinite training times does the weight matrix encode all PCs equally and recover the exact solution *W* = *I*. We thus see that, at any finite learning time, the output of the linear network will be more sensitive to the earlier PCs. Note that this non-uniformity in sensitivity emerges despite the lack of any constraints on *W* other than learning.

Having introduced this model to explain our findings in artificial networks, we next wondered how it would compare to human behavioral data. We first examined the sensitivity of the linear model to the spatial frequency of a sinusoidal grating (Fig. 5e). In adult humans, the detection threshold to changes in frequency increases linearly with frequency (Fig. 5g) (Caelli et al, 1983). To compare to human data, we can plot the “detection threshold” of an artificial network as the inverse squared sensitivity of the network output to frequency. This is proportional to the error rate of an optimal read-out of frequency given injected Gaussian noise (Rao, 1945). At several snapshots during training, we observed that the spatial frequency threshold increased linearly with frequency above a certain cutoff frequency, below which the threshold saturated at a low value (Fig. 5f). Even a single matrix trained to reconstruct images reproduces human-like sensitivity to spatial frequency when trained with gradient descent.

If the human perceptual system is also implicitly constrained by learning dynamics, this would be apparent in psychophysical studies of young children. Indeed, the highest observable frequency of a sinusoidal grating continues to improve with age even up to adolescence (Fig. 5i) (Leat et al, 2009; Mayer and Dobson, 1982). This is experience-dependent; when sight is restored in young children by the removal of cataracts their spatial acuity gradually improves (Maurer et al, 1999). These effects can be reproduced in our model of linear image reconstructions. By measuring the network’s spatial acuity as a function of training episode as the highest spatial frequency whose simulated detection threshold (inverse squared sensitivity) was below a fixed cutoff, we found we could reproduce a linear increase of spatial acuity with age (Fig. 5h). Learning with gradient descent reproduces not only an efficient encoding of spatial frequency but also the way in which visual acuity increases linearly with age.

The theory of learning in deep networks makes several further predictions for human perception, many of which have been explored previously (Saxe, 2015; Wenliang and Seitz, 2018). A central feature of this framework is a characteristic sigmoidal curve for perceptual learning tasks (Fig. 5c). Such sigmoidal learning curves are observable in humans on perceptual learning tasks that are sufficiently difficult (Fig. 5d) (Ahissar and Hochstein, 1997). This curve is sigmoidal because the rate of improvement depends upon the current level of sensitivity as well as the difference from asymptotic sensitivity (see Methods). This causes sensitivity to rise exponentially at first but eventually converge exponentially towards an asymptote. In human perceptual learning experiments, the learning curve is indeed better described as an exponential than other functional forms such as power laws (Dosher and Lu, 2007). Although gradient descent in linear systems is a simple model, it accurately captures the functional form of how perception improves with experience.

In the analysis above we trained towards the objective of reconstructing inputs, which is an unsupervised objective. However, the mathematical reason why gradient descent learns frequent inputs first also applies to supervised learning. For two features with equal correlation with the output labels but different variance in the inputs, the network will learn to use the higher-variance feature first (see Methods for derivation). Due to this additional bias beyond task usefulness, networks trained on a wide range of objectives will show greater sensitivity for frequent features.

The emergence of efficient coding in supervised tasks can be verified with a simple task in which the frequency and usefulness of input features are varied independently (Fig. 6). We trained a nonlinear 3-layer neural network to decode the orientation of a sinusoidal grating appearing with a set probability distribution. We also applied noise to the output labels to control the information in each stimulus about the labels. As expected, both the input frequency and output noise separately affect the sensitivity of learned representations (Fig. 6b,c). However, this could also be explained by the effect of frequency and noise on task usefulness, defined as the total information in the input dataset about each label. To demonstrate that gradient descent introduces an additional bias beyond task usefulness, we next adjusted the magnitude of the noise such that the total information is uniform across labels. This requires applying a greater level of noise onto the labels that are more common, balancing their effects on information. Even in this case, a higher sensitivity to input orientation emerges for more common orientations (Fig. 6d). This now provides a deeper intuition of our findings earlier that networks trained on object recognition are more sensitive to frequent features. The preference for frequent features is a general feature of learning with gradient descent and is separate from frequency’s effect on the information about labels.

**Fig. 6.**
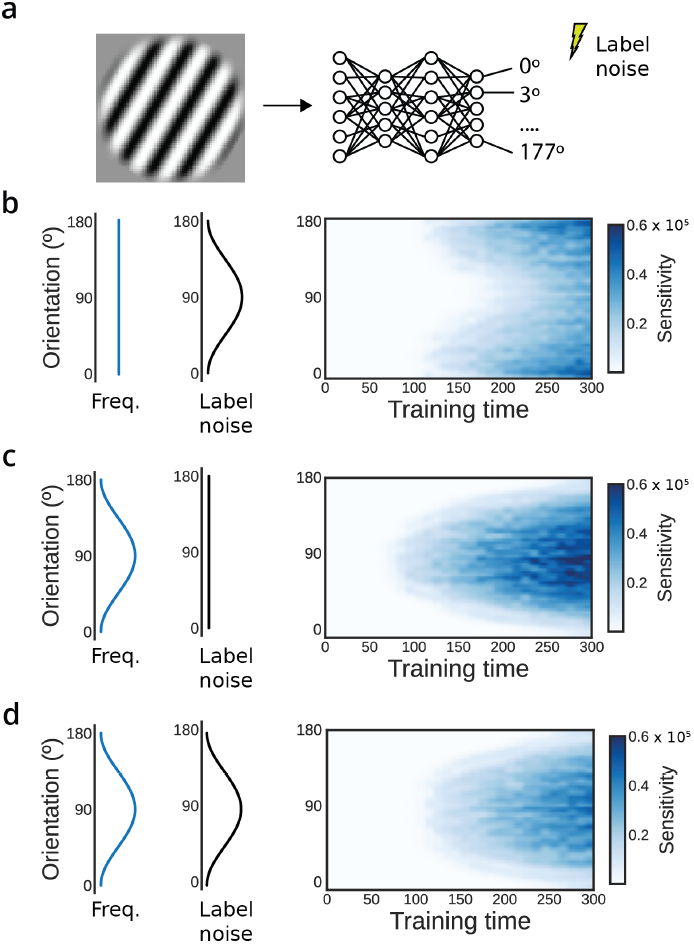
Dissociating the effect of frequency and information in supervised learning tasks. a) We trained 3-layer nonlinear neural networks to classify the orientation of sinusoidal gratings into 3º bins, varying either input frequency or output noise. b) We controlled the informativeness of input orientations by injecting noise into the labels as a function of orientation. The sensitivity of the first layer to input orientation is shown over learning. With uniform statistics, the more informative features are preferentially learned. c) The effect of varying input frequency without applying label noise. In this case, the more frequent features are preferentially learned. d) We then balanced noise and frequency such that the total information in the input dataset about each output label is uniform (see Methods). Learning with gradient descent still prefers common angles.

## Discussion

Here we found that the internal representations of artificial neural networks trained on ImageNet are more sensitive to basic visual features that are more common, which is a hallmark feature of efficient coding. We show that this hallmark naturally emerges from gradient-based learning. Even a minimal model of gradient-based learning – linear image reconstruction – reproduced human patterns of sensitivity to sensory variables and how these change over development. In this minimal model the dynamics of learning can be understood analytically. The correspondence of sensitivity and statistics emerges due to an implicit bias of gradient descent for common, high-variance aspects of the input data.

Our result provides a proof of principle that patterns of perceptual sensitivity in animals could be explained by a similar phenomenon. If plasticity in the brain approximates the gradient of some task, whether by reinforced Hebbian rules or some other algorithm, neural populations will preferentially encode the strongest dimensions in their inputs. As a result, organisms may not need dedicated learning algorithms for efficient codes in addition to algorithms for task-oriented learning. Many such local, unsupervised plasticity rules and neural control mechanisms have been previously proposed as ways the brain might develop efficient codes (Barlow, 1989; Bell and Sejnowski, 1995; Brito and Gerstner, 2016; Hyvärinen and Oja, 1997; Intrator and Cooper, 1992; Karklin and Simoncelli, 2011; Olshausen and Field, 1996; Ruderman and Bialek, 1993; Schwartz and Simoncelli, 2001; Zhou and Yu, 2018). Instead, any general algorithm approximating gradient descent may produce similar codes when learning towards many objectives.

It is important to note that our mathematical analysis of linear networks is highly simplified and may not accurately describe how learning affects sensitivity in general. A number of considerations complicate a generalization to nonlinear artificial neural networks, let alone brains. Nonlinearity makes linear decompositions inaccurate, and as a result we cannot say the precise features that gradient descent prefers to learn before others in nonlinear networks. New techniques from this emerging field may soon allow a more complete characterization of the dynamics of learning (e.g. Goldt et al (2020)). However, despite these caveats, we find that this model is useful to explain why efficient codes emerge in nonlinear artificial networks. It is remarkable that such a simple model of learning also captures qualitative features of human perceptual learning, as well. Learning in linear systems provides a valuable source of intuition for the effects of learning in more complex systems.

While we have shown one mechanism for how learning can induce a statistics/sensitivity correspondence, it is not the only mechanism by which it could do so. Theories of deep learning often distinguish between the “rich” (feature learning) and “lazy” (kernel) regimes possible in network learning (Woodworth et al, 2020). Our models reside in the rich regime, which involves learning new intermediate representations and assumes a small weight initialization. In the alternative, lazy regime, intermediate representations change little over learning and only a readout function is learned (Jacot et al, 2018). Interestingly, networks in the “lazy” (kernel) regime evolve under gradient descent as if they were linear in their parameters (Lee et al, 2019), and furthermore have the inductive bias of successively fitting higher modes of the input/output function as more data is presented (Bordelon et al, 2020; Canatar et al, 2021). The modes are defined differently, however, via the kernel similarity matrix rather than the direct input covariance. Despite the dissimilarities in these mathematical approaches, both center gradient descent and learn important aspects of the problem before others. These similarities suggest that a statistics/sensitivity correspondence could be derived for other network regimes.

Our broadest finding — that task-oriented learning can be a mechanism of producing efficient codes — is relevant to the discussion in psychophysics of the nature of the constraints implied by perceptual sensitivity patterns. It has long been recognized that these patterns imply some limitation upon coding capacity. Here, we make the distinction between implicit limitations due to (a lack of) learning and explicit limitations upon the maximum achievable code quality after learning, such as noise, metabolism, or a limited number of neurons. Although these categories limit perception with different mechanisms, they produce similar patterns of perceptual sensitivity. This distinction is particularly meaningful for explaining perceptual learning. Previously, perceptual improvements during development have been interpreted as a reduction in internal noise accompanied by a continuous maintenance of optimally efficient codes (Dosher and Lu, 1998; Kiorpes and Movshon, 1998). In our framework, learning naturally creates codes that reflect environmental statistics at all stages of learning, and there is no need to invoke a reduction in noise to explain improvements. This is consistent with the view that perceptual learning increases the signal-to-noise ratio through neuronal changes that enhance the signal strength (Gold et al, 1999; Schoups et al, 2001). To be sure, the nervous system is indeed constrained by hard ceilings such as noise and metabolism; learning probably ceases eventually. The implicit constraints due to learning are complementary to these and their relative contribution decreases with age and experience.

Although operating at a much shorter time-scale, sensory adaptation induces similar behavioral changes as perceptual learning, such as improving sensitivity to small stimulus differences at the adapted (i.e. learned) stimulus value. It has been argued that sensory adaptation is a form of efficient coding, optimally re-allocating sensory encoding resources according to recent stimulus history (Fairhall et al, 2001). While the adaptation induced changes in neural encoding characteristics such as reduction in response gain and changes in tuning curves are well characterized (Benucci et al, 2013; Kohn and Movshon, 2004), it is unknown how these local changes in neural representation accuracy depend on the specific details and dynamics of the sensory history. Thus it will be interesting to explore the degree to which sensory adaptation and its dynamics can be explained and predicted by the global objective of a task-dependent learning rule (gradient descent) in a continually updating (i.e., adapting) sensory processing system such as the brain.

The model system of gradient descent in linear systems can make several further predictions if taken as a model of human perceptual learning. In this model, the perceptual learning rate can be quantitatively modeled as a function of input statistics, importance, and current performance. These predictions could be verified in experiments that separably vary label noise and input statistics in supervised perceptual learning problems. Additionally, learning in the rich domain predicts that the learning system should represent the outside world in a low-dimensional way, with additional dimensions being added over time according to their variance and importance. As such, these learning dynamics naturally give rise to low-dimensional neural representations. Such learning dynamics may thus underlie the popular idea in neuroscience of low-dimensional neural manifolds (see Flesch et al (2022)).

A learning framework for perception points to a different sort of normative analysis of why we perceive the way that we do. Optimality can be defined in two ways. It can characterize the maximum achievable code quality, in an information-theoretic sense, given some number of neurons and their biological limitations. Alternatively, one might also describe responses that are optimal given the limited experience by which to learn the statistics of the world. Even ideal observers must learn from limited data, and successful learning from limited data must be constrained (Wolpert and Macready, 1997). Appropriate learning constraints would be selected for by evolution. Future research may help to unravel these optimal learning algorithms and characterize their sensory consequences.

## Methods

### Stimuli and calculation of network sensitivity

In all networks, we defined the sensitivity of a particular layer to a sensory variable as the squared magnitude of the gradient. For a layer with N nodes and vector of activations *y*, the sensitivity with respect to a sensory variable *θ* is:

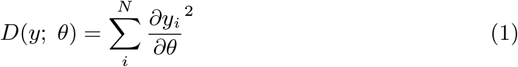

This definition of sensitivity can be related to the Fisher Information about a sensory variable *θ* in an artificial stimuli set. This is relevant for comparisons to human psychophysical data as the notion of sensitivity inferred from discrimination thresholds is the Fisher Information. In particular, our definition of sensitivity can be interpreted as the Fisher Information of systems with internal Gaussian noise of unit variance, and furthermore for the orientation of stimuli within an artificial stimulus ensemble with one stimulus per value of *θ*. A derivation of this connection can be found in the Mathematical Appendix.

The sensitivity can be calculated through backpropagation or by the method of finite differences. We created differentiable generators of stimuli in the automatic differentiation framework of Pytorch. This allowed calculating the sensitivity directly via in-built backpropagation methods.

For the figures in the text, we used Gabor stimuli with a spatial frequency of 2 cycles per 100 pixels, a contrast so that pixels span the range of [-1,1] in intensity in units of z-scored ImageNet image intensities, and a Gaussian envelope with *σ* = 0.5. We marginalized over the phase of the Gabor by averaging the sensitivity calculated with 10 linearly spaced spatial phases tiling the interval [*−π, π*]. The sinusoidal stimuli input to the linear network varied in spatial frequency, and we similarly averaged sensitivity over spatial phase. Finally, for the hue stimuli, we generated images of a uniform color in HSV color space and converted pixel values to RGB. Results were marginalized over the S and V axes in the range [.5, 1] which corresponds to the calculation of hue histograms on ImageNet (see below), which necessarily involves binning S and V.

### Deep nonlinear network experiments

To measure the sensitivity of pretrained networks, we first downloaded pretrained ResNet18 and VGG16 (with batch normalization) networks from the Torchvision python package (v0.11) distributed with Pytorch. For the vision transformer, we used a distribution in Python available at https://github.com/lukemelas/PyTorch-Pretrained-ViT. For each layer in these networks, we calculated sensitivity to orientation and hue were calculated with the stimuli generators described above. The ‘layers’ are defined differently for each network. For ResNet, layers are what in this architecture are called residual blocks (each of which contain multiple linear-nonlinear operations). For VGG, layers are the activations following each linear or pooling layer. Layers within the vision transformer are what are called transformer blocks.

We implemented a number of controls to determine the extent to which the observed patterns of sensitivity related to image statistics. We first ran the sensitivity analysis on untrained networks; we used the Pytorch default initialization. To ensure that the architectures do not show inherent patterns only in a certain regime of weight sizes, we calculated sensitivity on a copy of the networks in which the weights were shuffled. We wanted to preserve weight sizes in a layer-specific manner, and so shuffled the weights only within each tensor.

As further control on the effect of image statistics we retrained certain models on a version of ImageNet in which all images were rotated by 45º, or as well in which the hue of images were rotated by 90º. The transformer model was not retrained due to its expense. Image modifications were performed with Torchvision’s in-built rotation and hue adjusting image transformations.

We trained all networks using a standard training procedure: stochastic gradient descent with an initial learning rate of 0.1, decaying by a factor of 10 every 30 epochs, as well as a momentum value of 0.9, and a batch size of 256 images. The networks were trained for 90 epochs. To match the original training setup, we augmented the image dataset with random horizontal reflections and random crops of a size reduction factor varying from 0.08 to 1. Note that the random horizontal reflections change the statistics of orientations so as to be symmetric around the vertical axis. After training the sensitivity was calculated as above.

### ImageNet hue statistics

We wrote a custom script to extract the hue histogram of all pixels in all images in the ImageNet training set. We binned hues with a resolution of 1º, and binned hues over the S and V range [.5,1] to focus on strongly colored pixels. The exact range is arbitrary, but importantly matches the range used when calculating network sensitivity.

### Linear network experiments

We first constructed a database of 32x32 images of natural scene image portions. These image portions were extracted from ImageNet (Deng et al, 2009), made greyscale, and cropped to size. Our constructed dataset contained over 100,000 examples of image portions. We then performed PCA on this dataset using the PCA method in Scikit-Learn (Pedregosa et al, 2011), and displayed the singular values of the top 1,000 components in Figure 5.

Our task consisted of reconstructing these image portions using a single- or multilayer fully-connected linear neural network. To ensure no architectural bottleneck exists, the internal (hidden) dimension of the multilayer network remained at 32^2^, the same as the input and output. The initial parameter values of the networks were scaled down by a factor of 100 from the default Pytorch initialization to ensure rich-regime learning. Networks were trained to minimize the mean-squared error of reconstruction using stochastic gradient descent, a learning rate of 1.0, and a batch size of 16,384, the largest that would fit in memory. The large batch size minimizes effects relating to batch stochasticity.

During learning, we calculated the sensitivity to spatial frequency as well as the projection of the learned weight matrix upon the PCA basis vectors of the inputs. The projection upon each PCA vector is given by 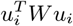, where *W* is the product matrix corresponding to the linear network and *u*_*i*_ is the *i*th PCA component. The sensitivity to spatial frequency was calculated by constructing a sinusoidal plane wave test stimulus with parameterized frequency and phase and using Pytorch’s automatic differentiation capability to obtain the derivative of network output with respect to frequency. The sensitivity was calculated for 64 equally-spaced phase offsets and the result averaged over phase.

### Supervised label noise experiment (Fig. 6)

In this experiment we trained a 3-layer neural network with ReLU nonlinearities to decode the orientation of 64x64 pixel image of a sinusoidal grating. The period of the sinusoid was 12.8 pixels, and in each stimulus the sinusoid carried a random phase offset. The random phase and orientation ensured that no image was repeated. In each image the orientation was sampled in the interval [0, *π*] from a specified probability distribution (either a uniform distribution or 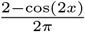. The objective was the categorization of images into 60 bins of orientations, with success quantified via a cross-entropy loss function.

The addition of noise to the output labels was calibrated such that, on average over a dataset, any orientation *θ* is as informative as any other despite a potentially nonuniform orientation distribution *p*(*θ*). Since the total information in a dataset about a (potentially noised) label *y*_*θ*_ scales linearly with how often it appears, all else held equal, the variation in per-example information must exactly balance the change in frequency. That is, for any two orientations *θ*_*i*_ and *θ*_*j*_ and their corresponding (noised) labels 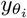 and 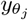, it must be that 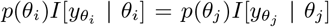 Here *I*[·] represents the information gained about a label having observed an input, i.e. the change in entropy over *y*_*θ*_ from the uniform distribution. This proportionality is satisfied if 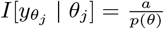 for some constant *a*.

Our approach thus requires applying label noise of a known entropy that varies with orientation. Because we optimize a cross-entropy objective, rather than e.g. a mean-squared-error objective, there are no interactions between neighboring bins. We applied noise by treating the nonzero element of the each label vector, which are indicator (1-hot) vectors, as a Bernoulli variable with rate *σ*(*θ*). *σ* = 1 corresponds to the zero-noise condition, and with rate 1 − *σ*(*θ*), a label is dropped out. For this noise, the information about each label is *I*[*y*_*θ*_ | *x*_*θ*_] = 1 − *H*_*b*_(*σ*(*θ*)), where *H*_*b*_(*σ*) is the binary entropy function. Together, the rate of Bernoulli noise is given by 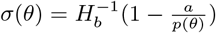. We approximated the inverse binary entropy function with a table lookup and assuming *σ ≥* 0.5.

### Sensitivity analysis of linear networks

In this section we will analyze the sensitivity of a linear multilayer neural network in which the weights of layer *i* are parameterized by *W*_*i*_. The output of such a network with N layers is:

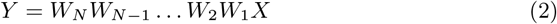

The product matrix is simply *W*, such that *Y* = *WX*.

Throughout this analysis we will make heavy use of the singular value decomposition of the weight matrix, which defines matrices *U*, *S*, and *V* such that *W* = *USV* ^*T*^. The matrix *S* is diagonal, and the diagonal elements are called the singular values *σ*_*i*_. The *U* and *V* matrices are orthonormal.

Our analysis describes how learning dynamics in this system acts to link output sensitivity to input statistics. Note that the derivation here is for arbitrary objectives; the instance of a reconstruction loss discussed in the main text is a special case. The analysis is organized in three stages: 1) how the sensitivity depends on the singular values *σ*_*i*_ of *W*, 2) how *σ*_*i*_ change with learning, and 3) how *σ*_*i*_ correspond to the image statistics.

### Sensitivity depends on the singular values of *W*

We wish to derive the sensitivity of a linear network to arbitrary input features. We will first examine the case of determining the sensitivity of the network the following feature: how much the data aligns with each *j*th singular vector of *W*. This is a weight-dependent feature. Specifically, let the feature *θ*_*j*_ be the dot product of the data with the *j*th right singular vector of the weight product matrix, 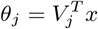. This feature is important as it can be used to analyze the sensitivity to arbitrary features.

For this feature, we find the sensitivity of the network is 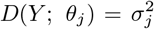. This result is intuitive, as 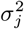 describes how much data lying along the vector *v*_*j*_ is amplified when multiplied with *W*. A derivation can be found in the Appendix. Thus, when 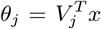, the sensitivity *D*(*Y* ; *θ*_*j*_) is constant and is the square of the associated singular value.

The sensitivity to more general *θ* can be understood using this result. This is because the key derivative can be decomposed into the derivatives with respect to right singular vectors: 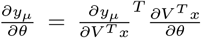. In this case, we find that 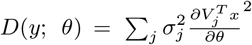 (see Appendix for derivation). Thus, for arbitrary *θ*, the sensitivity depends on how 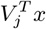 depends on *θ* times the size of the associated singular value, summed over components *j*.

### The behavior of the singular values

Previous literature describes how the weight matrix changes due to gradient descent (Arora et al, 2019; Saxe et al, 2013). More information about these results can be found in the Appendix.

We first define a (potentially data-dependent) cost function:

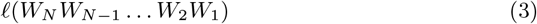

As described by Arora et al (2019), under certain conditions on the weight initialization the direction of the unit vectors *u* and *v* rotate with learning in a specific way. Note that they remain unit length during learning. This result, quoted in the Mathematical Appendix as Theorem 2, states that the vectors are static when they align with the singular vectors of the gradient of the loss, ∇*ℓ*(*W*). More specifically, if the singular vectors are static then *U*^*T*^ ∇*ℓ*(*W*)*V* is diagonal. This will become an important condition for tying the input statistics to the singular values of *W*.

Another important result from previous literature describes how singular values of the product matrix *W* evolve as a function of time *t*:

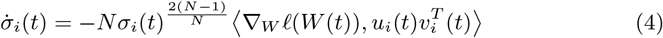

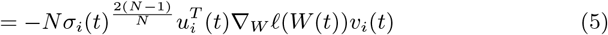

Thus, each singular value evolves as a product of a function its current size and the network depth 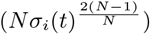 multiplied by how much the gradient correlates with the corresponding singular vectors. This formalism assumes continuous learning dynamics; see (Gidel et al, 2019) for a treatment of finite step sizes.

### Relation of frequency to sensitivity

In this section we wish to show how the input statistics affect the singular vectors and values of *W*. Our approach is to show that frequency *p*(*θ*) reflects in the covariance of *θ*. The covariance affects the rate of learning of the singular values of the weight matrix *W*.

### Frequency vs. variance

In our analysis of how the statistics of data affect sensitivity, we focus on the variance of features. Since previous literature in psychophysics focus on frequency as defined by the vector *p*(*θ*) with a scalar value for each orientation *p*(*θ* = *θ*_*j*_) (e.g. Wei and Stocker (2015)), it is appropriate to discuss their relation. Our analysis focuses on variance in part because attributes like orientation measured can occur with a real-valued strength in each image patch when measured by e.g. Gabor filters or Fourier decomposition. Thus a description of *p*(*θ*) in natural images might be more completely characterized with a two-dimensional matrix with dimensions for angle and intensity. Variance summarizes the intensity axis and characterizes how unpredictable each orientation is within each image patch. The second reason we work with variance is that it cleanly relates to the speed of learning.

When features are binary and either present or not, variance and frequency are closely related. Modeling presence as a Bernoulli variable, the frequency is the probability *p*(*θ*_*j*_) and the variance is *σ*(*θ*_*j*_) = *p*(*θ*_*j*_)(1 − *p*(*θ*_*j*_)). Note that at very small values of *p*(*θ*_*j*_), *σ*(*θ*_*j*_) *∼ p*(*θ*_*j*_). However, features that are nearly always present (*p*(*θ*_*j*_) near 1) have a very low variance. It is interesting to note that this behavior aligns with the expectation that efficient sensory systems should dedicate more resources to features whose presence is uncertain (*p*(*θ*_*j*_) = .5) than to those whose presence is guaranteed (*p*(*θ*_*j*_) = 1). Variance is thus very similar to absolute frequency for rare Bernoulli variables and in general may be a more intuitive measure of feature importance in regards to what determines efficient patterns of sensitivity.

### What W learns: autoencoding objective

Further describing the growth of singular values requires a choice of objective. The base case of our study is the autoencoding objective defined for a set of inputs *X*:

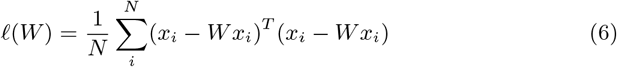

Our goal is to determine how *W* evolves for this cost function. We will examine both the singular vectors and the singular values.

During learning, the singular vectors rotate (recall they are unit length and orthogonal) until they reach a fixed point. For this cost function, it is easy to verify that a fixed point of dynamics is when the singular vectors are equal to the principal components of the inputs (see Appendix for proof). That is, the vectors are static when Σ_*xx*_ = *V* Λ*V* ^*T*^ and *W* = *V SV* ^*T*^ for the same *V* but potentially different Λ and *S*. This alignment is especially relevant given the expression for network sensitivity derived above. With the vectors aligned, the sensitivity to each corresponding principal component of the inputs is given by 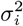, the squared singular value of W.

The evolution of sensitivity is thus governed by the evolution of singular values. The rate of change of *σ*_*i*_ is complicated to calculate because the singular vectors can potentially rotate. However, for the sake of analysis one can examine the case when the singular vectors are initialized at the fixed point mentioned above, as in previous literature (Saxe et al, 2013). In this set of initial conditions, the time-evolution of each singular value of W is given by (Saxe et al, 2013; Arora et al, 2019):

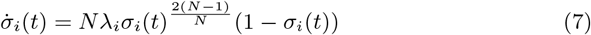

Note that the rate of learning is controlled by *λ*_*i*_, the standard deviation of the *i*th principal component of the inputs. The term on the right causes *σ*_*i*_(*t*) to converge to 1 asymptotically, as is expected as the solution of the full-rank reconstruction problem is *W* = *I*. For deeper networks (*N ≥* 2), the growth is sigmoidal and approaches a step function as *N → ∞* (see Gidel et al (2019)). Thus, in this axis-aligned initialization, the singular values *σ*_*i*_(*t*) are learned in order of the variance of the associated principal components of the inputs.

Together, these results mean that the sensitivity of a linear network’s output to the principal components of the inputs evolve in order of variance when trained on input reconstruction. This is exactly the case for the axis-aligned initialization and approximately true for small initializations. For the single-matrix network displayed in the figure in the main text, the sensitivity to the *j*th PC thus evolves over time *t* as:

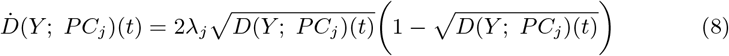

### What W learns: supervised learning

We can also determine how input statistics affect the sensitivity for the more general class of objective functions when *Wx* is trained to match some target *y* by minimizing the mean-squared error:

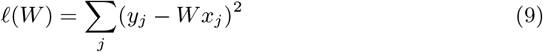

As before, we can gain intuition about W by beginning from an initialization that is axis-aligned with the final solution. For the supervised case, these initializations share the singular vectors of the data/labels, but can differ in the singular values. Given Σ_*xx*_ = *V* Λ*V* ^*T*^ and Σ_*xy*_ = *UT V* ^*T*^, we set *W* (0) = *USV* ^*T*^ for the same *U* and *V*. See the Appendix for proof that this is a fixed point of singular vector dynamics.

This initialization allows us to understand how the singular values of the weight matrix change. As derived in the Appendix, the time evolution of *σ*_*i*_ is given by:

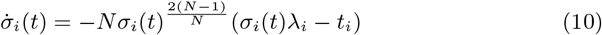

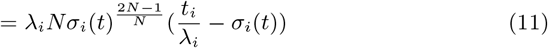

As in the case for input reconstruction, the *i*th singular value approaches a target. Instead of 1, this value is 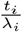, the ratio of the importance of this component (the input/output singular value *t*_*i*_) and the standard deviation of that component in the inputs *λ*_*i*_. The growth rate is controlled by the distance from this asymptote (right term) and as well as on the input statistics *λ*_*i*_. Thus, even for the case of supervised learning the input statistics affect what is learned first via gradient descent directly through Σ_*xx*_ via *λ*_*i*_, and not just through the input/label covariance Σ_*xy*_.

## Code availability

All code used to create the figures is available at https://github.com/KordingLab/ANN_psychophysics.

## Acknowledgements

The authors are appreciative to research assistance in earlier versions of this project from Ryan Guan and Ryan Jeong.

## Supplemental figures

**SI Fig 1:**
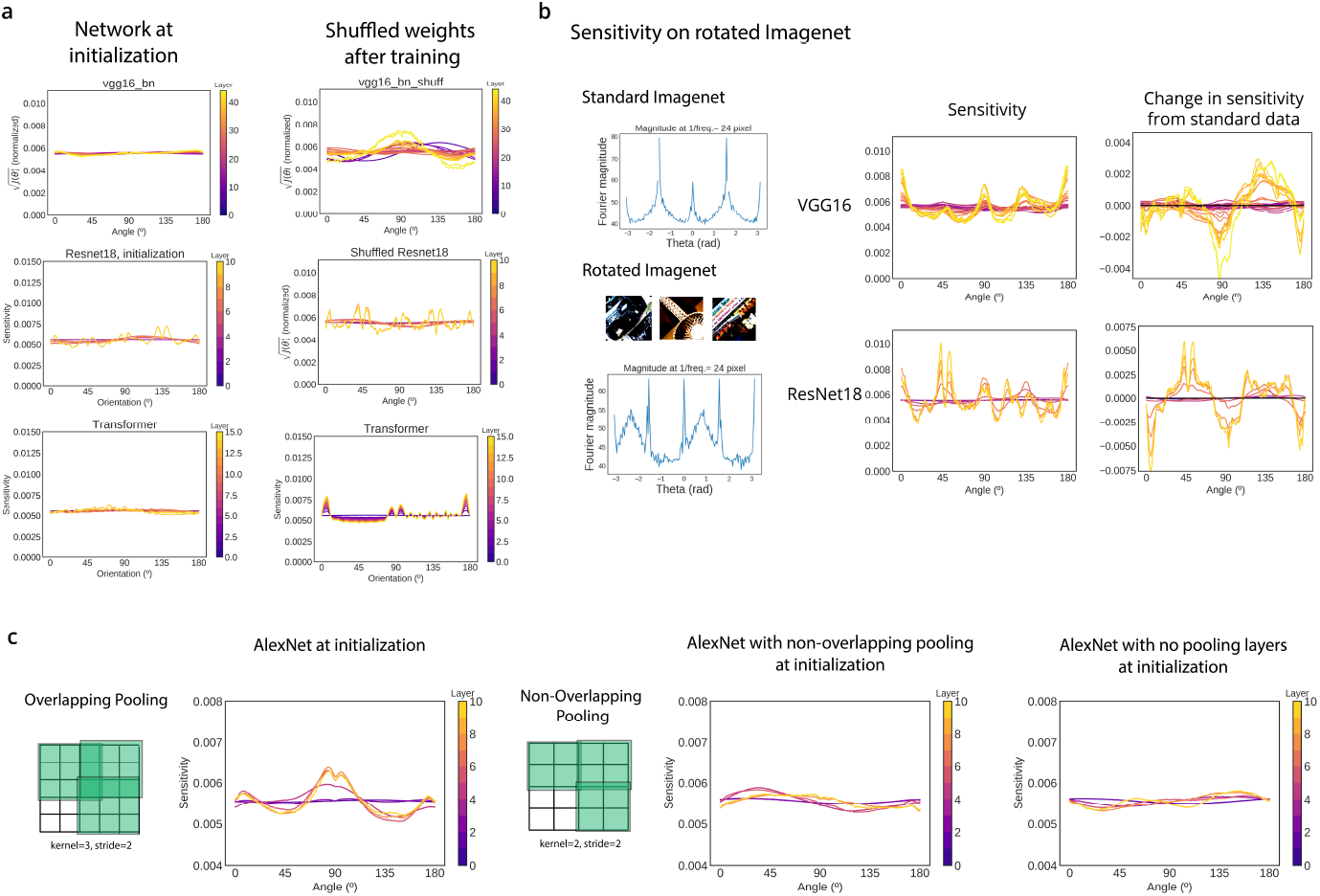
Controls for orientation sensitivity analyses. a) The sensitivity to the orientation of Gabor stimuli (see Methods for stimuli parameters) at initialization, left column, and after shuffling the parameters within each layer, right column. b) We retrained the Resnet18 and VGG16 architectures on a version of ImageNet in which all images were rotated by 45º. Left: The orientation statistics can be observed as the magnitude of the Fourier spatial decomposition around a circle centered at the origin in frequency space. This method of analysis will show artifacts of spikes at the cardinals due to the edge effects of rectangular images, but it is a useful control that image statistics do change with rotation. Top is for standard ImageNet, and bottom is for rotated ImageNet. Middle column: The sensitivity of ResNet18 and VGG16 after retraining on rotated images. Though changed, it does not appear as a simple shift of the patterns seen for standard ImageNet (Fig. 2). Right: The difference of this sensitivity pattern from the sensitivity pattern observed after training on standard ImageNet shows that changes do correspond to the change in image statistics, at least in part. c) One source of training-independent sensitivity to the cardinal orientations is overlapping pooling, where it is used. (None of the three above networks employ overlapping pooling.) Left: In AlexNet, which does, the network shows a strong non-uniformity of sensitivity at initialization. Right: The use of non-overlapping pooling greatly diminishes the non-uniformity, as does the complete removal of pooling layers (and accompanying change in the convolutional filter downsampling).

**SI Fig. 2:**
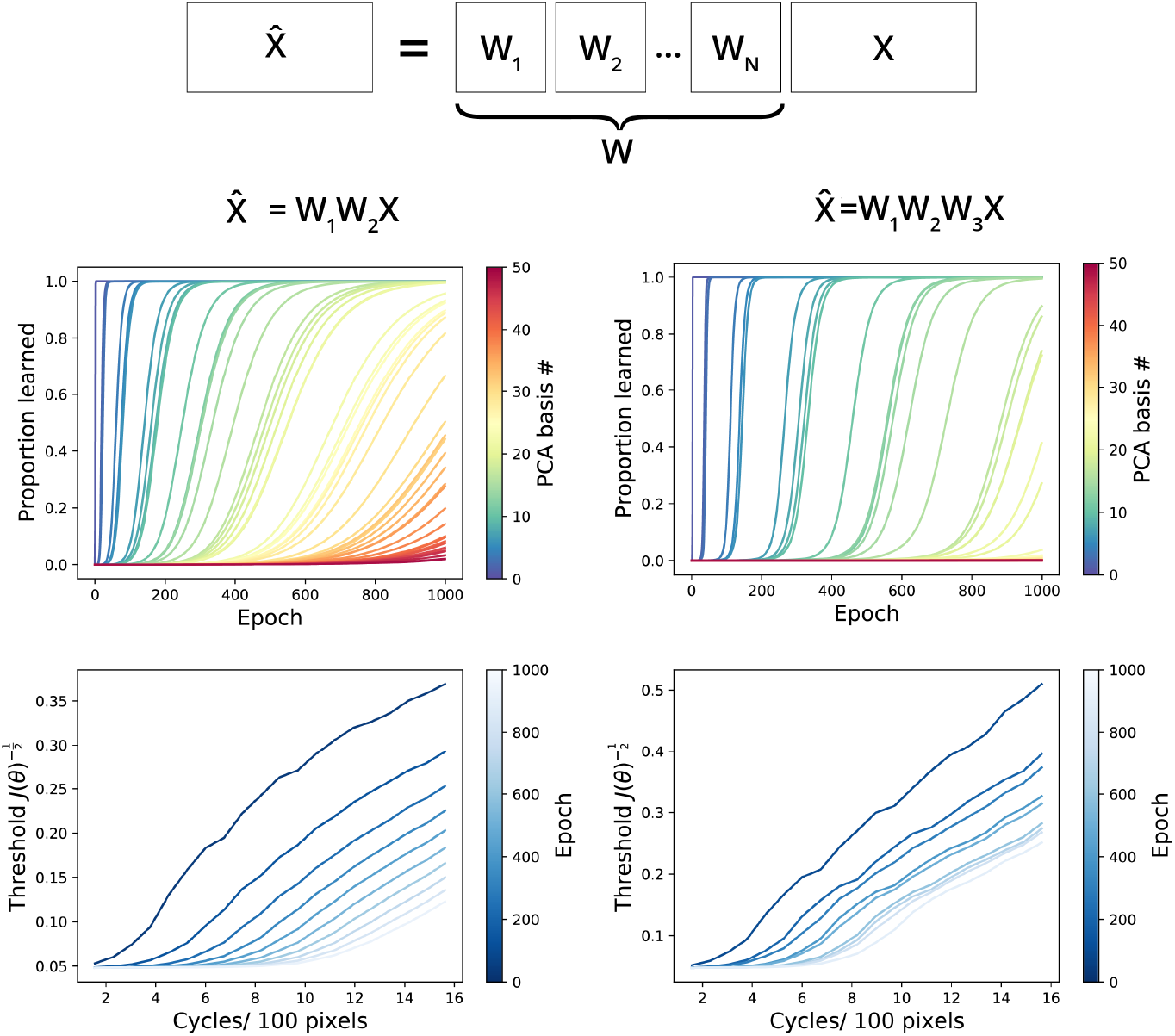
Learning dynamics of 2 and 3 layer linear networks. Top: As the depth of networks increases, the principal components are learned in sharper transitions. Bottom: The “threshold” of spatial frequency detection, defined as the inverse square root of the sensitivity to frequency, shows similar patterns regardless of network depth.

## Appendix

### Relation of sensitivity to the Fisher information

In the main manuscript, we define the *sensitivity* of a data point *y* to a feature *θ* as:

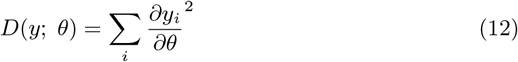

Throughout this Appendix, lower-case variables represent vectors and capital letters refer to matrices. We denote the sensitivity an ensemble of data points as *D*(*Y* ; *θ*).

#### The Fisher Information

In this section we show that the above definition of sensitivity can be interpreted as the Fisher Information of *y* given *θ* for the case of Gaussian internal noise of unit variance.

Suppose after obtaining *y*, we observe *noisy* observations of it 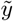. Then, the Fisher information of 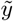 about some input feature *θ* is

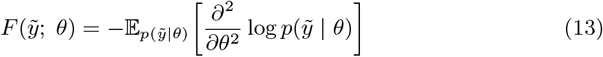

When *y* is a representation of some inputs *x*, the Fisher information is that implicitly reflects the stimulus ensemble *X*, through the connection *x → θ → y*. It is often the case that we wish to measure the *average* Fisher information over such an input ensemble. For example, the Fisher Information of orientation on natural images, or on test stimuli. We denote such an average by with capital letters, suggesting a matrix of examples: 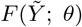.

##### Fisher information for internal noise sources

We are concerned with the Fisher information 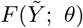 when only a single input *x* corresponds to each *θ*. For example, one could be interested in the Fisher information about orientation as characterized by a set of rotating sinusoidal gratings with identical contrast, phase, and frequency.

Since we observe *Y* noisily, we still observe a distribution 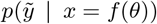 for each *θ*. Here we have written *x* = *f* (*θ*) for some real function *f* : ℝ *→* ℝ since *x* is deterministic given *θ*. In this case the Fisher information is

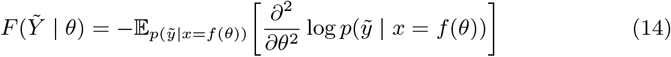

This is the expression we focus on in this paper. It is simple to calculate using derivatives (as detailed below) and a valid point of comparison to human data characterized with a stimulus ensemble with only one *θ* per *x*.

##### The addition of external noise sources

As an aside, if one wants to determine the Fisher information about orientation *on natural images*, the desired quantity changes. In naturalistic datasets there are many inputs *x* that could correspond to each value of *θ*. Thus, there are two sources of noise: *external* noise producing the distribution *p*(*x* | *θ*), and *internal* noise producing 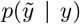 or alternatively 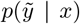. In this case the Fisher information is not easily obtained from derivatives of model representations. This is because this quantity requires marginalizing over *x* inside of the Fisher expression:

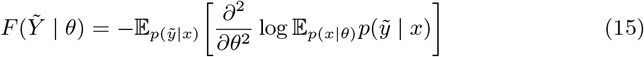

The expectation inside of the log makes this analytically intractable. Approximations may be useful for rather simple *p*(*x* | *θ*), but this is prohibitive for naturalistic data. Thus our expression for sensitivity is not comparable to the Fisher information about data with external noise.

#### Gaussian internal noise

The notion of network sensitivity in our paper can be equated with Fisher Information in the case that one observes a noised 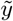 containing an additive injection of zero-mean Gaussian noise. Thus with noise *ζ ∼ 𝒩* (0, 1),

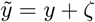

In this case 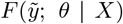 simplifies. Since the noise is independent over outputs *y*_*i*_ and furthermore Gaussian over each output unit,

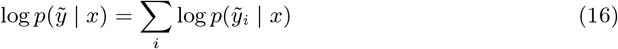

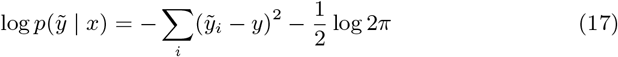

Taking the derivative and expectation over *ζ*, we obtain the well-known result for the Fisher of Gaussians:

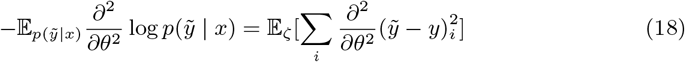

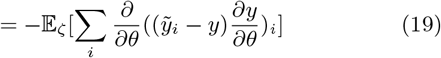

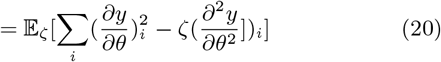

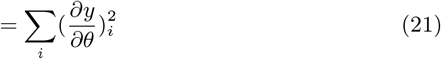

#### Sensitivity analysis of a linear network

Imagine that we have a linear multilayer neural network in which the weights of layer *i* are parameterized by *W*_*i*_. The output of such a network with N layers is:

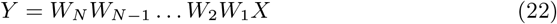

The product matrix is simply *W*, and *Y* = *WX*.

The total sensitivity can be broken up into terms that depend on the decomposition of *W*. This will be the bridge to a theory of learning.

As a minimal example, let us first examine the case of determining the sensitivity as a function of how much the data aligns with each singular vector of *W*. That is, our feature *θ*_*j*_ is the dot product of the data with the *j*th right singular vector of the weight product matrix. As discussed later, for autoencoding cost functions this will align with the principal components of the data after a bit of training, and so this might also be said to be the sensitivity about the *j*th principal component.

Defining the SVD and feature of interest as,

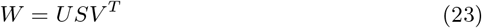

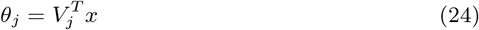

For this feature, the expected sensitivity for an input ensemble *p*(*x*) is:

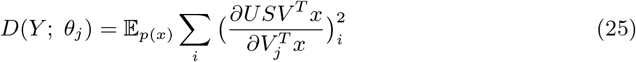

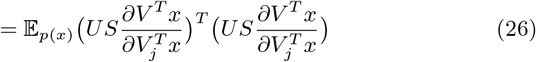

Since 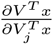 is a one-hot vector that is 1 in the *j*th row and zero everywhere else, it acts to “pick out” the *j*th column of *US*. Note that the expectation over *p*(*x*) disappears as well.

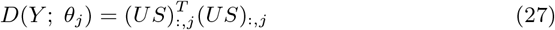

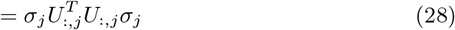

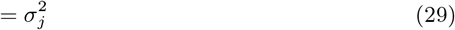

Here 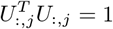 because columns of U are orthonormal. Thus, for when 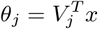 the sensitivity *D*(*Y* ; *θ*)_*j*_ is constant and is the square of the associated singular value.

#### More general *θ*

More general *θ* can be understood using the result of the last section. The approach is to decompose an arbitrary derivative into the derivatives with respect to right singular vectors, as such: 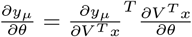. In this case,

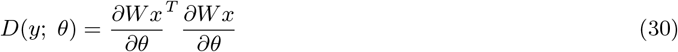

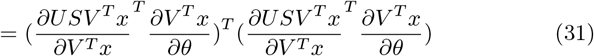

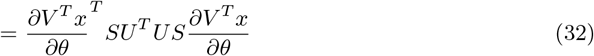

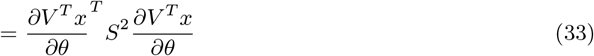

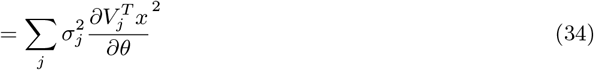

Thus, for arbitrary *θ*, the sensitivity depends on the derivative of the *j*th right singular vector with respect to *θ* times the size of its associated singular value.

### The behavior of the singular values

Let us now establish a cost function upon the product matrix.

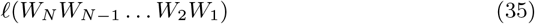

### Results of previous literature

Though many papers have adopted this framework, as cited in the main text, here we quote the result of Arora et al (2019).

***Summary:***

*During gradient descent, under certain restrictive conditions on the initial values of W*_*i*_, *the* ***singular values*** *of the product matrix evolve qualitatively differently for N* = 1 *vs. N >* 1. *For N >* 1 *they grow larger sigmoidally (roughly one-at-a-time) and in order of their contribution to the cost ℓ*(*W*).

The results to follow examine what happens when we train *W*_*i*_ via gradient descent to minimize *ℓ*(*W*). Each *W*_*i*_ now becomes a function of time, *W*_*i*_(*t*), and

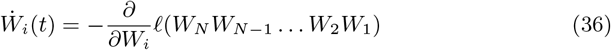

In addition we assume that the matrices are initialized in a *balanced* manner, meaning that for all *j < N*,

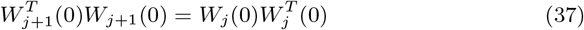

This holds approximately when the weights are initialized very close to zero.

***Lemma (Arora et al. 2019)***

*The product matrix W(t) can be expressed as:*

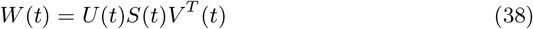

*where U* (*t*) *and V* (*t*) *have orthonormal columns and S*(*t*) *is diagonal*

Our theory hinges on the behavior of the diagonal elements of *S*(*t*), which we will denote as *σ*_*i*_(*t*).

#### Theorem 1 (Arora et al. 2019)

*The singular values of the product matrix W (t) evolve by:*

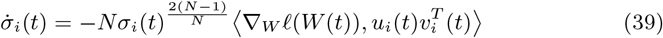

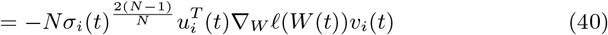

Thus, each singular value evolves as a product of a function its current size and the network depth 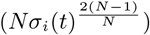 multiplied by how much the gradient correlates with the rank-1 matrix implied by the singular vectors. Note that if *N* = 1 there is no dependence on the current size of *σ*_*i*_(*t*).

Another important result concerns the rotation of the unit vectors *u*(*t*) and *v*(*t*). It states that the vectors are static when they align with the singular vectors of ∇*ℓW* (*t*).

#### Theorem 2 (Arora et al. 2019)

*Assume that at initialization, the singular values of the product matrix W (t) are distinct from zero, and that the matrix factorization is non-degenerate, i*.*e. has depth N ≥* 2. *Then, for any time t such that the singular vectors of the product matrix W (t) are stationary, i*.*e*. 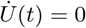 *and* 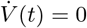, *then U*^*T*^ (*t*)∇*ℓ*(*W* (*t*))*V* (*t*) *is diagonal*.

### Relation of input statistics to sensitivity

Our approach is to show that frequency *p*(*θ*) reflects in the covariance of *θ*. This in turn affects the rate of learning of the singular values of the weight matrix *W*, at least for certain objectives. This connects the sensitivity, or Fisher Information of 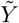, back to the frequency.

#### The data covariance affects the learning of *σ*_*i*_(*t*): autoencoding objective

The base case of our study is the autoencoding objective defined for a set of inputs *X*:

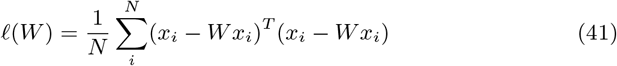

Our goal is to determine the evolution of *σ*_*i*_(*t*) that results from this cost function. First, see that

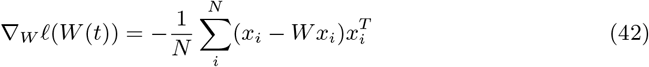

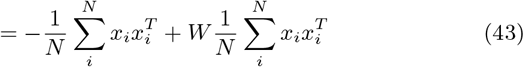

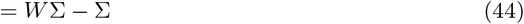

Here Σ is the data covariance, assuming *X* is centered.

In general, the time evolution of each singular value *σ*_*i*_(*t*) is complicated to calculate because the singular vectors can potentially rotate, i.e. 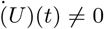. However, for the sake of analysis we can examine a limited case when the direction of the singular vectors is static. This will allow us to obtain an analytic expression for the evolution of the singular values in terms of the data covariance. In particular we will examine the case when the weight matrix is initialized to share right singular vectors (but not singular values) with the data covariance.

By plugging the expression for ∇_*W*_ *ℓ*(*W* (*t*)) into Theorem 2, it can be seen that if Σ = *V* Λ*V* ^*T*^ and *W* = *V SV* ^*T*^ for the same *V*, then

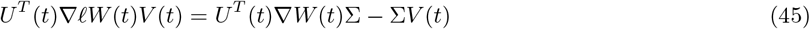

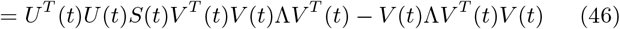

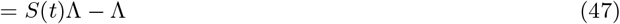

This is diagonal, and thus by Theorem 2 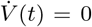 during gradient descent on the autoencoding objective.

By Theorem 1, this initialization results in:

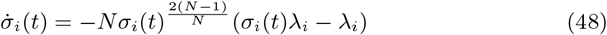

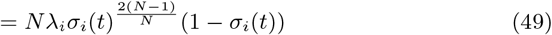

#### Extension to supervised learning

A more general class of objective functions is when *Wx* is trained to match some target *y*. The input statistics are again relevant here. If we again take the mean-squared error as the objective,

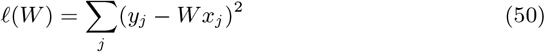

The evolution of the singular values is determined by the gradient,

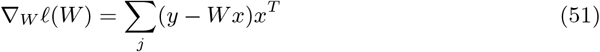

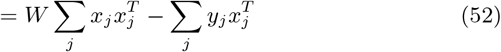

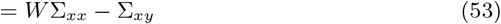

By Theorem 1, then, we have that,

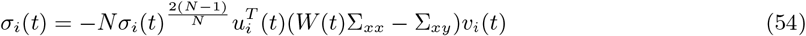

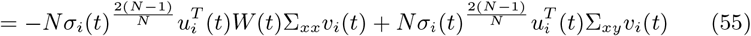

Thus, the evolution of the singular values depends on two additive terms. One of these (left) has no dependence on the labels *y*, only on the statistics of the data.

As before, we can gain intuition about this evolution by beginning from an initialization that is axis-aligned with the final solution. For the supervised case, these initializations share the singular vectors of the data/labels, but can differ in the singular values. Given Σ_*xx*_ = *V* Λ*V* ^*T*^ and Σ_*xy*_ = *UT V* ^*T*^, we set *W* (0) = *USV* ^*T*^ for the same *U* and *V*. This means that,

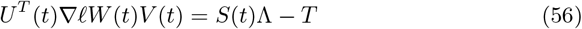

This is a diagonal matrix, and thus a fixed point of learning.

For this initialization, then,

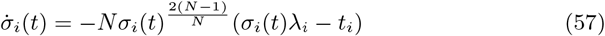

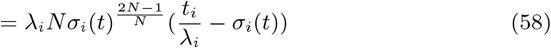

## Notes

### Competing Interest Statement

The authors have declared no competing interest.

